# Differences in functional guild responses to elevation and topography in Himalayan ant communities

**DOI:** 10.64898/2026.06.11.731772

**Authors:** Netra Kadambi, Mansi Mungee, Ramana Athreya

**Affiliations:** Indian Institute of Science Education & Research Pune, India; Azim Premji University, Bhopal, India

**Keywords:** Ants, Functional Diversity, Multi-guild, Elevational profile, Microclimate, Eastern Himalayas

## Abstract

We investigated the influence of small-scale topographic and associated microclimatic drivers on the elevational patterns of functional alpha and beta diversity in ant communities of eastern Himalayas. Ants were sampled along three elevational transects spanning ∼1000–2000 m on two nearby mountain slopes, but differing in aspect, inclination and summit height. Ant genera were quantified at the quadrat level, yielding an abundance of 3090 genus-quadrats (from 22577 individuals in 1215 quadrats), almost evenly split between the two mountain slopes. Each ant was assigned to guilds along functional axes of thermal niche, trophic position and competitive abilities. Comparative analyses of functional diversity profiles along the two mountain slopes revealed no differences in the composite multi-guild alpha and beta diversity but significant differences amongst individual guild profiles. Seven of the eleven functional guilds showed strong elevational trends along the steeper north-facing slope, compared to only two of eleven on the shallower south-facing slope. These results indicate that similar functional space can be maintained despite taxonomic turnover within guilds or shifts in their relative abundances highlighting the influence of fine-scale topographic variation on community assembly. Alternatively, these findings demonstrate that the pattern of multivariate functional diversity across elevation does not necessarily provide an understanding of changing composition of the communities therein.

## Introduction

Evaluating the contributions of various factors that simultaneously drive the diversity and composition of communities has been a central theme of many ecological studies (Anderson et al., 2011). Montane ecosystems serve as natural laboratories to investigate the role of environmental and geographic factors governing spatiotemporal patterns of biodiversity (Perrigo et al., 2019; Rahbek et al., 2019). Pronounced climatic and resource heterogeneity across elevational gradients promote high regional diversity through rapid turnover of communities (McCain & Grytnes, 2010; Sundqvist et al., 2013). The complex topography of montane systems also generates environmental heterogeneity, creating a matrix of microhabitats which influences community assembly (Rahbek et al., 2019; Stein et al., 2014). At the landscape level, topographic features such as ridges and valleys may also influence connectivity between local communities by creating barriers to and corridors for dispersal (Perrigo et al., 2019).

Many studies have examined the effect of large-scale macroclimatic drivers, such as temperature, in shaping communities. However, fewer studies have investigated the effect of local topographic features which modify environmental conditions at finer spatial scales (König et al., 2024; Potter et al., 2013). Factors such as aspect and inclination can affect the diversity and function of communities through differences in local conditions dictated by solar radiation, wind exposure, temperature-moisture balance and soil nutrient availability (Moeslund et al., 2013). Hence, communities from the same regional species pool may assemble differently on neighbouring slopes with similar elevational extents.

Functional diversity patterns complement the information from taxonomic patterns in understanding the ecological mechanisms shaping community composition (Swenson, 2011). As such, functional traits of individuals, and not their taxonomic identity, interact with the biotic and abiotic components of the environment, consequently shaping community composition (Kraft et al., 2015). Multiple studies have quantified functional diversity by sampling morphological, physiological, phenological or behavioural traits that are “functional,” i.e. those that relate to an ecosystem service (Funk et al., 2017; Violle et al., 2007). The number, type and resolution of traits is subjective, depending on the taxonomic group, ease of measurement and resource constraints (Dawson et al., 2021; Jansen et al., 2018). Also, the traits sampled may not necessarily be “functional” in the local environmental context, introducing noise to our inferences of the underlying processes structuring communities (Bellwood et al., 2019; Mlambo, 2014; Violle et al., 2007)

As an alternative, studies have examined responses of discrete functional types or groups, which may represent a cluster of correlated, rather than individual, functional traits (Blaum et al., 2011). In some cases, this approach has been shown to reflect community processes better than linking individual traits to their hypothesised ecological functions (Raffard et al., 2017). Moreover, given the paucity of functional trait data across many faunal groups, the functional guild framework offers a more pragmatic approach to study community assembly (Rubio & Swenson, 2024). While useful, this approach ignores intragroup variation, which can substantially influence community structure (Mungee et al., 2021; Palacio et al., 2025; Spasojevic & Suding, 2012; Violle et al., 2012).

We investigated the functional guild composition of ant communities on two mountain slopes in the eastern Himalayas of Arunachal Pradesh, in north-east India. Ants (Family: Formicidae) are an excellent system to examine functional responses of communities along environmental gradients. Although belonging to a single family, ants exhibit remarkable ecological diversity, occupying multiple trophic levels (Drager et al., 2023), and a wide range of ecological niches (Uemori et al., 2022) (Drager et al., 2023). Their distributions are shaped by both abiotic factors, such as environmental conditions and resource availability, and biotic interactions, particularly competition, which strongly influences patterns of coexistence (Davidson, 1980; but also see Cerda et al., 2014). This sensitivity to environmental and ecological filters has made ants a focal taxon in functional ecology which has also led to the development of the Global Ants Database in 2006 (Gibb et al., 2017) and numerous studies examining functional community structure across environmental gradients. However, despite this global interest, functional information on ant communities from the Indian subcontinent remains sparse. Only two datasets are currently represented in the Global Ants Database from the Indian subcontinent: the Western Ghats (Basu, 1997) and the western Himalayas (Bharti et al., 2009). Bharti et al. (2013) proposed a generic-level functional guild classification for Himalayan ants, providing a practical framework for examining functional diversity and community assembly in these underrepresented communities.

Eastern Himalayas are one of the world’s most biodiverse regions in the world (Myers et al., 2000). The steep slopes expose communities to strong climatic gradients over relatively short distances, allowing the strong effects of environmental changes to be examined with reduced influence from dispersal and historical biogeography. The complex topography of these young fold mountains generates substantial local variation in slope aspect, inclination, and ruggedness resulting in distinct microclimates even among neighbouring slopes experiencing the same regional climate (Feldmeier et al., 2020; König et al., 2024; Geres et al., 2025). This provides an excellent opportunity to disentangle the influence on diversity patterns of local microclimatic variations from those of broader macroclimatic gradients. Ground-dwelling arthropods, particularly ants, are well suited for examining these processes because their distributions are strongly influenced by temperature, moisture, and soil conditions, all of which can vary markedly over fine spatial scales in montane environments (Choi & Jang, 2022; Gilgado et al., 2022; Theron et al., 2022).

In this study, we ask whether local topographic variation can modify elevational patterns of functional diversity within the same regional landscape. Specifically, we examine how functional diversity in ant communities changes between two nearby mountain slopes that differ in topography and associated microclimatic conditions. We used available information on thermal niches, trophic roles, and competitive strategies of ant genera to investigate if the elevational profiles of functional guilds differed between the slopes. We predicted that neighbouring slopes would exhibit different functional alpha and beta diversity profiles despite experiencing the same broad climatic gradient, reflecting differences in local environmental filtering.

## Methods

### Study site

Ants were sampled along three elevational transects on two mountain slopes in West Kameng district of Arunachal Pradesh during May−August 2021 **(Figure 1).** We had one transect spanning 900─1850 m (geographic distance of 5.6 km) inside Eaglenest Wildlife Sanctuary (27°02‘–09’N, 92°18‘–35’E) and two transects near Khupi model village (27°11‘–27°12’N, 92°38‘–39’E) spanning 1050–2050 m (geographic distances of 2.9 and 3.6 km). The Khupi transects are north-facing, terminate close to the ridge at 2100 m, have a highway cutting across them at ∼1650 m, and are separated from each other by 0.1-1.8 km along contours. On the other hand, the Eaglenest transect is south-facing, terminates well below the ridge at 3250 m, and has little or no anthropogenic disturbance. Eaglenest and Khupi are separated by 30 km of aerial distance and ∼100 km along elevational contours (30-metre satellite data).

**Figure 1:**
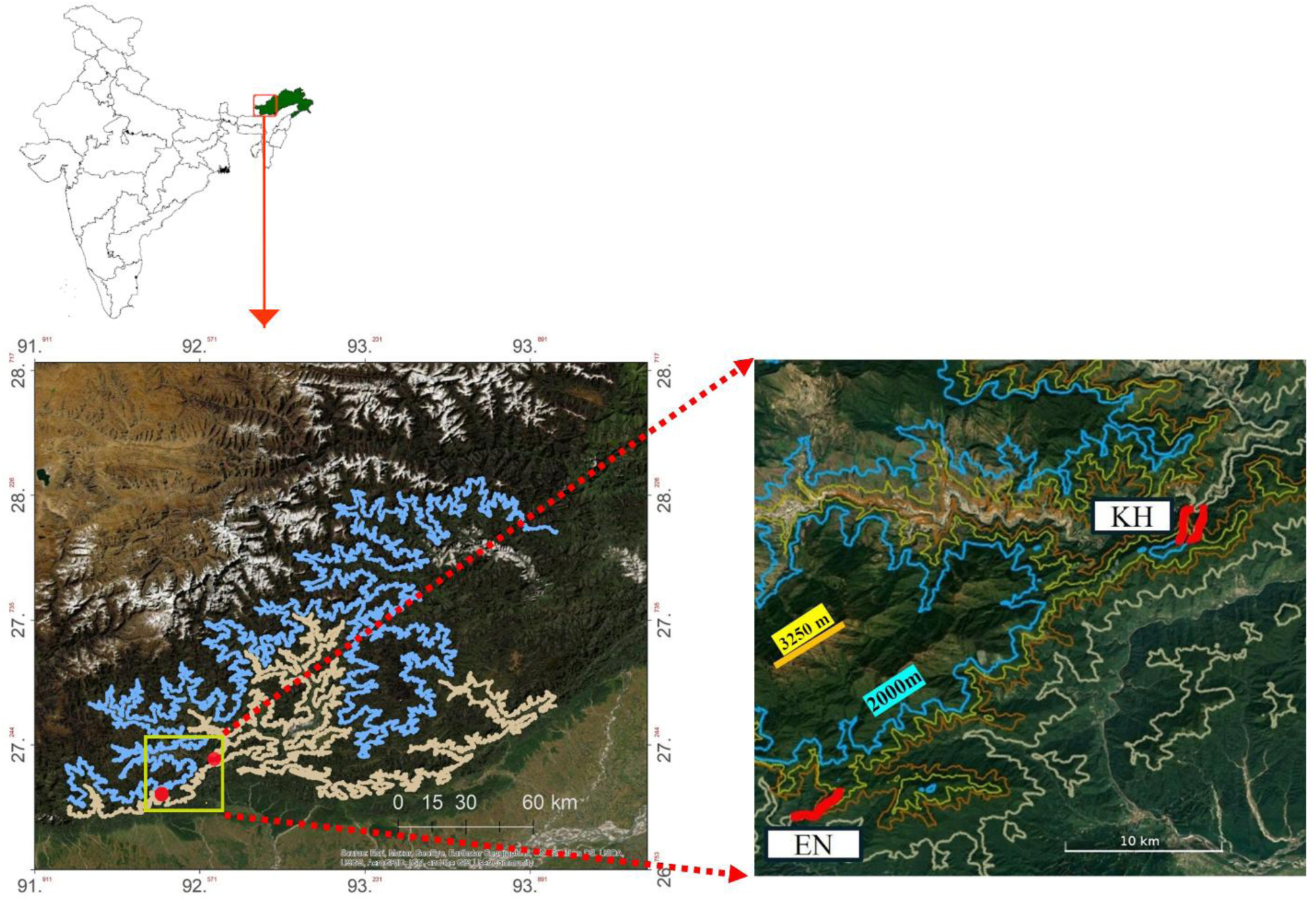
Contour maps of Western Arunachal Pradesh showing the two sites, Eaglenest Wildlife Sanctuary (EN) and Khupi Model village (KH), where ant communities were sampled. The map on the left shows the 1000 m (cream) and 2000 m (blue) contour line with the two sites represented as red solid circles inside the yellow box. The map on the right shows four contour lines spaced 250 m apart in elevation (1000 m a.s.l.– cream, 1250 m a.s.l.– brown, 1750 m a.s.l.– mustard, 2000 m a.s.l. – blue). The thick red line represents the three elevational transects: one in Eaglenest (EN: 950 m–1850 m a.s.l.) and two transects at Khupi (KH1: 1150–2100 m a.s.l.; KH2: 1050–1950 m a.s.l.). The yellow bar represents the ridge at Eaglenest. Note that the transects at Khupi (KH1 and KH2) terminate at the local ridge (2100 m a.s.l.), represented by the blue circular contour line (2000 m a.s.l.).

### Sampling strategy

We used Winkler traps (Agosti et al., 2000) to sample ground dwelling ants from quadrats (1m x 1m) along the three transects. Sampling nodes were spaced 30 m apart in ground distance. At each node, ants were sampled in three quadrats located 10 m apart along a line approximately perpendicular to the transect. In total, we sampled 564 quadrats in Eaglenest and 651 in Khupi (combining both transects). We estimated generic abundances by counting the number of quadrats in which a genus occurred, to avoid inflated abundance due to aggregation of foraging workers from the same nest. We segregated the quadrats in Eaglenest and Khupi into elevational communities of about 100 m width and approximately equal abundance **(Figure 2)**.

**Figure 2:**
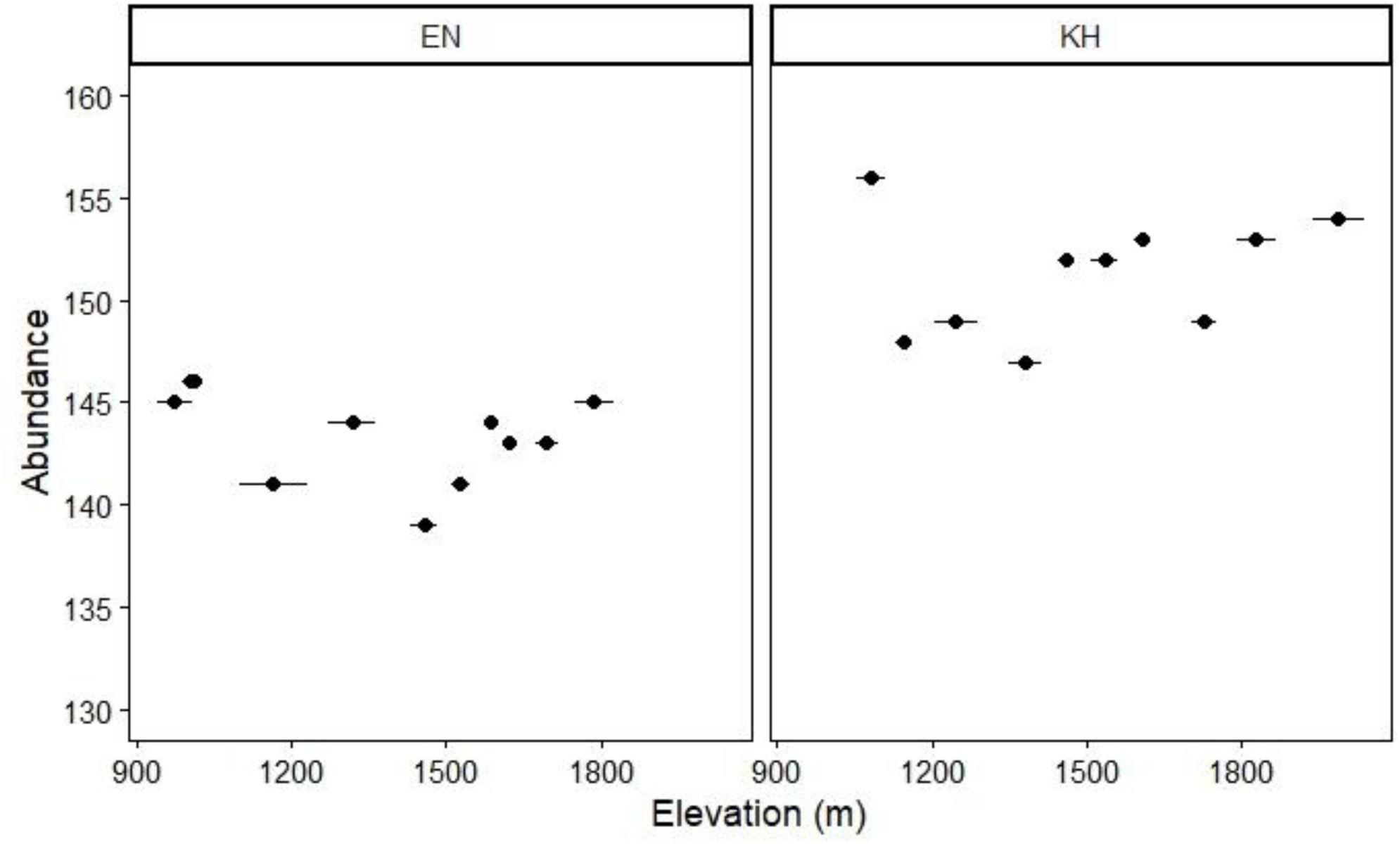
Ant elevational communities in Eaglenest (EN = 11 communities) and Khupi (KH = 10 communities). Generic abundance for each community is plotted against the elevation (mean ± sd) and are represented by solid black circles for both Eaglenest and Khupi. Communities were obtained by segregating sampling nodes to obtain roughly equal abundances of communities for a given transect. On average, communities are spaced approximately 80–100 m apart in elevation. Note that abundance of elevational communities in Eaglenest and Khupi range between 139–146 and 147–156, respectively.

### Functional guild framework in ants

**Table 1** shows the assignment of all ant genera to one guild in each of three functional axes, viz. (i) thermal niche (abiotic axis) (ii) trophic level (biotic and abiotic axes) and competitive dominance (biotic axis) following available information from AntWiki and the functional classification scheme for Himalayan ant genera (Bharti et al., 2013). The guilds within the three functional axes are listed below:

**Table 1:** Genus-level assignment of binary scores along three functional axes (thermal niche, trophic level and competitive dominance) examined. Each genus is assigned into mutually exclusive guilds within an axis. TCS = tropical climate specialists, CCS = cold climate specialists, TG = thermal generalists; Herb = herbivore, Omni = omnivore, SP = small predator, LP = large predator; SL = specialists/competition avoiders, OP. = opportunists, SB = subordinates, DM = dominants.

| Genus |  | Thermal Niche |  |  | Trophic Level |  |  |  | Competitive Dominance |  |  |  |
| --- | --- | --- | --- | --- | --- | --- | --- | --- | --- | --- | --- | --- |
|  |  | TCS | CCS | TG | HB | OM | SP | LP | DM | SB | OP | SL |
| 1 | *Anochetus* | 1 | 0 | 0 | 0 | 0 | 0 | 1 | 0 | 0 | 0 | 1 |
| 2 | *Aphaenogaster* | 0 | 0 | 1 | 0 | 1 | 0 | 0 | 0 | 0 | 1 | 0 |
| 3 | *Brachyponera* | 1 | 0 | 0 | 0 | 1 | 0 | 0 | 0 | 0 | 1 | 0 |
| 4 | *Camponotus* | 0 | 0 | 1 | 0 | 1 | 0 | 0 | 0 | 1 | 0 | 0 |
| 5 | *Carebara* | 1 | 0 | 0 | 0 | 0 | 1 | 0 | 1 | 0 | 0 | 0 |
| 6 | *Cataulacus* | 1 | 0 | 0 | 0 | 1 | 0 | 0 | 0 | 0 | 0 | 1 |
| 7 | *Cerapachys* | 1 | 0 | 0 | 0 | 0 | 0 | 1 | 0 | 0 | 0 | 1 |
| 8 | *Crematogaster* | 0 | 0 | 1 | 0 | 1 | 0 | 0 | 1 | 0 | 0 | 0 |
| 9 | *Diacamma* | 1 | 0 | 0 | 0 | 0 | 0 | 1 | 0 | 0 | 0 | 1 |
| 10 | *Discothyrea* | 1 | 0 | 0 | 0 | 0 | 1 | 0 | 0 | 0 | 0 | 1 |
| 11 | *Dolichoderus* | 0 | 0 | 1 | 0 | 1 | 0 | 0 | 0 | 0 | 1 | 0 |
| 12 | *Ectomomyrmex* | 1 | 0 | 0 | 0 | 0 | 0 | 1 | 0 | 0 | 0 | 1 |
| 13 | *Emeryopone* | 1 | 0 | 0 | 0 | 0 | 1 | 0 | 0 | 0 | 0 | 1 |
| 14 | *Hypoponera* | 1 | 0 | 0 | 0 | 0 | 1 | 0 | 0 | 0 | 0 | 1 |
| 15 | *Lasius* | 0 | 1 | 0 | 0 | 1 | 0 | 0 | 0 | 1 | 0 | 0 |
| 16 | *Leptogenys* | 1 | 0 | 0 | 0 | 0 | 0 | 1 | 0 | 0 | 0 | 1 |
| 17 | *Lophomyrmex* | 1 | 0 | 0 | 0 | 1 | 0 | 0 | 0 | 0 | 0 | 1 |
| 18 | *Monomorium* | 0 | 0 | 1 | 0 | 1 | 0 | 0 | 0 | 0 | 1 | 0 |
| 19 | *Myopopone* | 1 | 0 | 0 | 0 | 0 | 0 | 1 | 0 | 0 | 0 | 1 |
| 20 | *Myrmecina* | 0 | 0 | 1 | 0 | 0 | 1 | 0 | 0 | 0 | 0 | 1 |
| 21 | *Myrmoteras* | 1 | 0 | 0 | 0 | 0 | 0 | 1 | 0 | 0 | 0 | 1 |
| 22 | *Nylanderia* | 1 | 0 | 0 | 0 | 1 | 0 | 0 | 0 | 0 | 1 | 0 |
| 23 | *Odontomachus* | 1 | 0 | 0 | 0 | 0 | 0 | 1 | 0 | 0 | 0 | 1 |
| 24 | *Ooceraea* | 1 | 0 | 0 | 0 | 0 | 1 | 0 | 0 | 0 | 0 | 1 |
| 25 | *Pheidole* | 0 | 0 | 1 | 0 | 1 | 0 | 0 | 1 | 0 | 0 | 0 |
| 26 | *Polyrhachis* | 1 | 0 | 0 | 0 | 1 | 0 | 0 | 0 | 1 | 0 | 0 |
| 27 | *Ponera* | 0 | 0 | 1 | 0 | 0 | 1 | 0 | 0 | 0 | 0 | 1 |
| 28 | *Pristomyrmex* | 1 | 0 | 0 | 0 | 0 | 1 | 0 | 0 | 0 | 1 | 0 |
| 29 | *Proceratium* | 1 | 0 | 0 | 0 | 0 | 1 | 0 | 0 | 0 | 0 | 1 |
| 30 | *Pseudolasius* | 1 | 0 | 0 | 1 | 0 | 0 | 0 | 0 | 0 | 0 | 1 |
| 31 | *Stictoponera* | 1 | 0 | 0 | 0 | 0 | 0 | 1 | 0 | 0 | 0 | 1 |
| 32 | *Strumigenys* | 1 | 0 | 0 | 0 | 0 | 1 | 0 | 0 | 0 | 0 | 1 |
| 33 | *Stigmatomma* | 1 | 0 | 0 | 0 | 0 | 1 | 0 | 0 | 0 | 0 | 1 |
| 34 | *Technomyrmex* | 1 | 0 | 0 | 0 | 1 | 0 | 0 | 0 | 0 | 1 | 0 |
| 35 | *Temnothorax* | 0 | 1 | 0 | 0 | 1 | 0 | 0 | 0 | 0 | 0 | 1 |
| 36 | *Tetramorium* | 0 | 0 | 1 | 0 | 0 | 1 | 0 | 0 | 0 | 1 | 0 |
| 37 | *Vollenhovia* | 1 | 0 | 0 | 0 | 0 | 1 | 0 | 0 | 0 | 0 | 1 |
| 38 | *Vombisidris* | 1 | 0 | 0 | 1 | 0 | 0 | 0 | 0 | 0 | 0 | 1 |

**Functional axis 1: Thermal niche**

1. Tropical-climate specialists (TCS): Genera that are largely in moist and warm tropical regions.
2. Cold-climate specialists (CCS): Genera that are largely in cold temperate regions.
3. Thermal generalists (TG): Genera adapted to both warm and cold climates with widespread distribution in both tropics and temperate regions.

**Functional axis 2: Trophic level**

1. Herbivores (HB): Genera that feed on sugary sap, seed or plant material.
2. Omnivores (OM): Genera that feed on both plant material and other arthropods as active predators or detritivores.
3. Small predators (SP): Cryptic predatory ants that hunt in small subterranean crevices and in layers of leaf litter. They also tend to be small in body size.
4. Large predators (LP): Large-bodied predatory ants that are unable to access small crevices and primarily hunt on more planar surfaces.

**Functional axis 3: Competitive dominance**

1. Dominant (DM): Behaviourally aggressive and numerically superior genera that directly outcompete other genera via interference competition.
2. Subordinate (SB): Genera co-existing with dominant genera by employing submissive strategies, such as nesting in suboptimal sites, maintaining smaller colonies or foraging in smaller territories.
3. Opportunistic (OP): Genera that exhibit flexible ecological strategies, thriving in disturbed and stressful habitats.
4. Specialists (SL): Genera that avoid competition by separating their nesting or foraging niche.

### Data analyses

We estimated the relative abundance of all guilds within each functional axis in each location, and for each elevational community. We used the Wilson confidence intervals for bi-/multinomial distributions to estimate the errors on the relative abundances. We excluded one unidentified genus (N = 21) and the genus *Mayriella* (N = 4) from the analyses due to lack of guild information.

We measured Gower’s functional distance (Gower, 1971) between each generic pair by assigning binary values (different:1, same:0) in each of the three functional axes. These values were averaged over the three axes, yielding functional distances of 0, 0.33, 0.66 or 1 for any pair. Note that the guilds within an axis are mutually exclusive, nominal and equidistant variables.

Rao’s quadratic entropy was used to represent the functional α-diversity of each elevational community. It was calculated using the *dbFD* function in the R package ‘*FD’* v1.0.12.3 (Laliberté et al., 2009). We fitted linear models to examine the association between elevation and functional α-diversity and relative abundance profiles of individual guilds.

We examined the abundance-weighted composite (multi-guild) functional β-diversity (FβD) with increasing elevational distance between communities. Functional β-diversity was estimated using the R package *‘BAT’* v.2.11.0 (Cardoso et al., 2015). The total functional dissimilarity (FβD) was partitioned into turnover and nestedness components (Baselga, 2013).

We also investigated the elevational profiles of functional beta diversity for individual guilds. In this we selected all genera in a community belonging to a particular guild (in one axis) and analysed the beta diversity of their occupancy of the guilds in the other two axes.

We used linear instead of the usual exponential growth/decay model for functional β because the estimated growth factor was very small (∼10^−5^ per km of elevational distance). We used the transect identity (Eaglenest v/s Khupi) as an interaction term in the linear models to test for transect-specific differences in the elevational responses.

## Results

We recorded a total of 22577 ants spanning 40 genera and 8 subfamilies, yielding 3,090 genus-quadrat (abundance) records. They were evenly split between Eaglenest (1577) and Khupi (1513). Of these, 3065 records from 38 genera were used for functional diversity analyses.

We observed significant differences in the relative abundances of the individual guilds **(Figure 3)**. Cold-climate specialists were rare in both Khupi and Eaglenest. Tropical-climate specialists dominated over thermal generalists in Eaglenest but were of similar abundance in Khupi. The rank hierarchy of guilds along the trophic level axis was conserved between Eaglenest and Khupi, despite substantial differences in relative abundances of some guilds. Small (leaf-litter) predatory ants were the most abundant guild followed by omnivores and larger (surface-foraging) predatory ants. Herbivores formed a negligibly small fraction in both Eaglenest and Khupi. However, large predators were more numerous in Eaglenest (LP = 20–25 %) than in Khupi (LP = 7–10 %).

**Figure 3:**
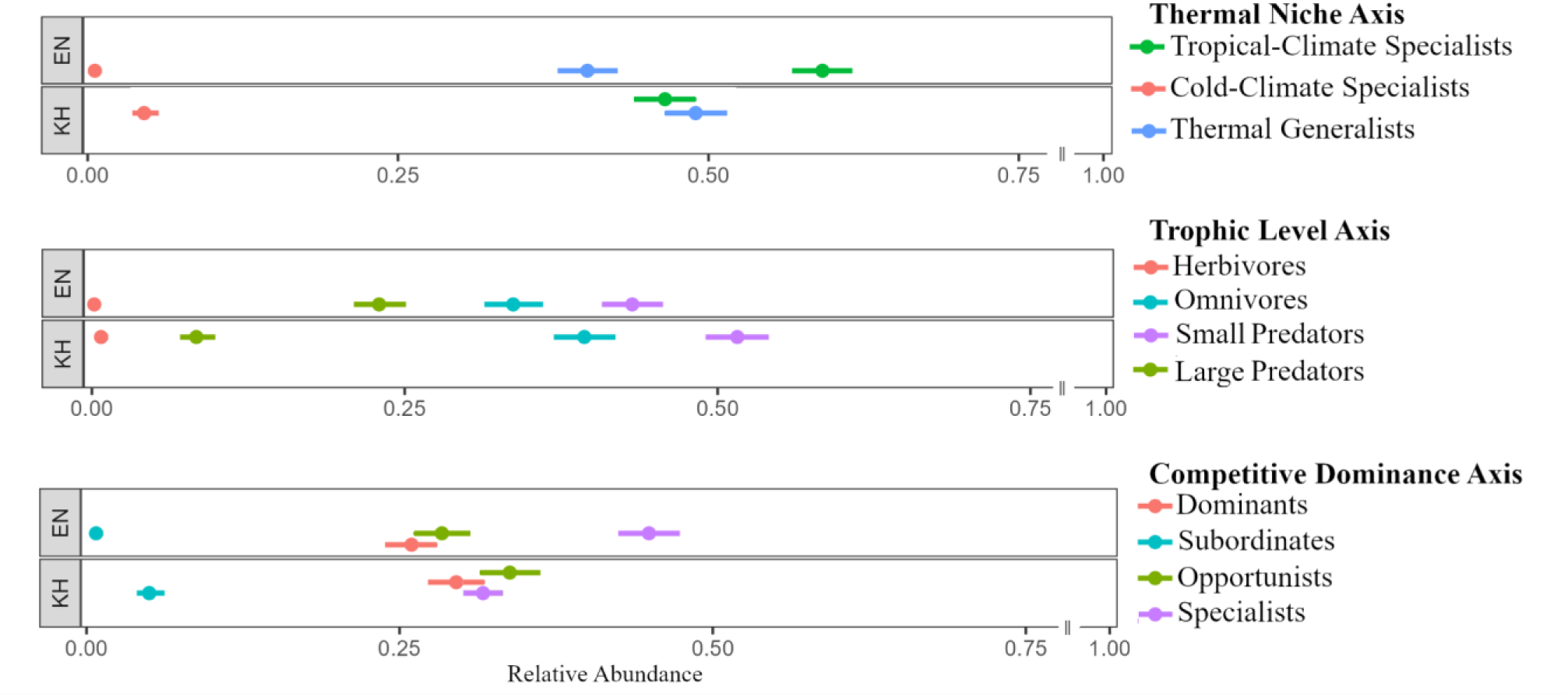
Mean relative abundance of ant guilds in each of the three functional axes in Eaglenest and Khupi. The three panels plot the distribution of proportions of each guild in each of the three functional axes (Thermal Niche, Trophic Level and Competitive Dominance), with each panel consisting of values in Eaglenest (EN) at the top and Khupi (KH) at the bottom. Mean values are obtained from a binomial distribution and plotted alongside Wilson’s binomial confidence intervals (95%). The identity of the eleven guilds is provided on the right under each functional axis.

Elevational profiles of relative abundances for the ant guilds are shown in **Figures 4-6**. The parameters of the fit are listed in **Table 2**.

**Figure 4:**
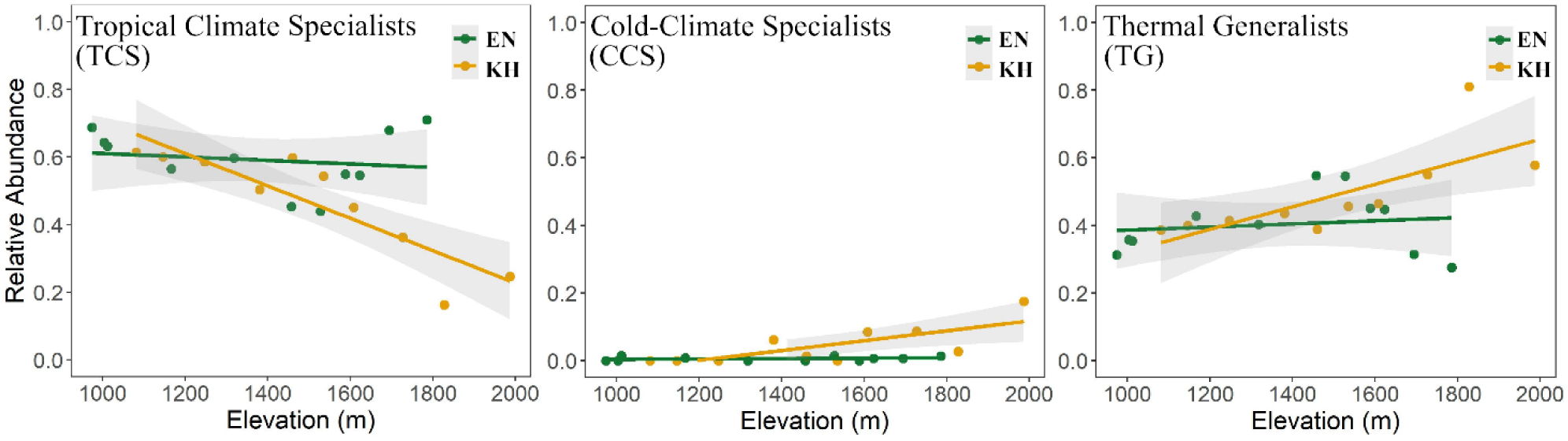
Relative abundance profiles of ant thermal guilds in Eaglenest (EN) and Khupi (KH) along 1000–2000 m span. Each green and yellow solid circle represent an elevational community in Eaglenest and Khupi, respectively. Green and yellow trend lines are the linear fits showing the magnitude of change in the proportion of each thermal guild with elevation. Abbreviations: TCS = tropical climate specialists, CCS = cold climate specialists, TG = thermal generalists.

**Table 2:**
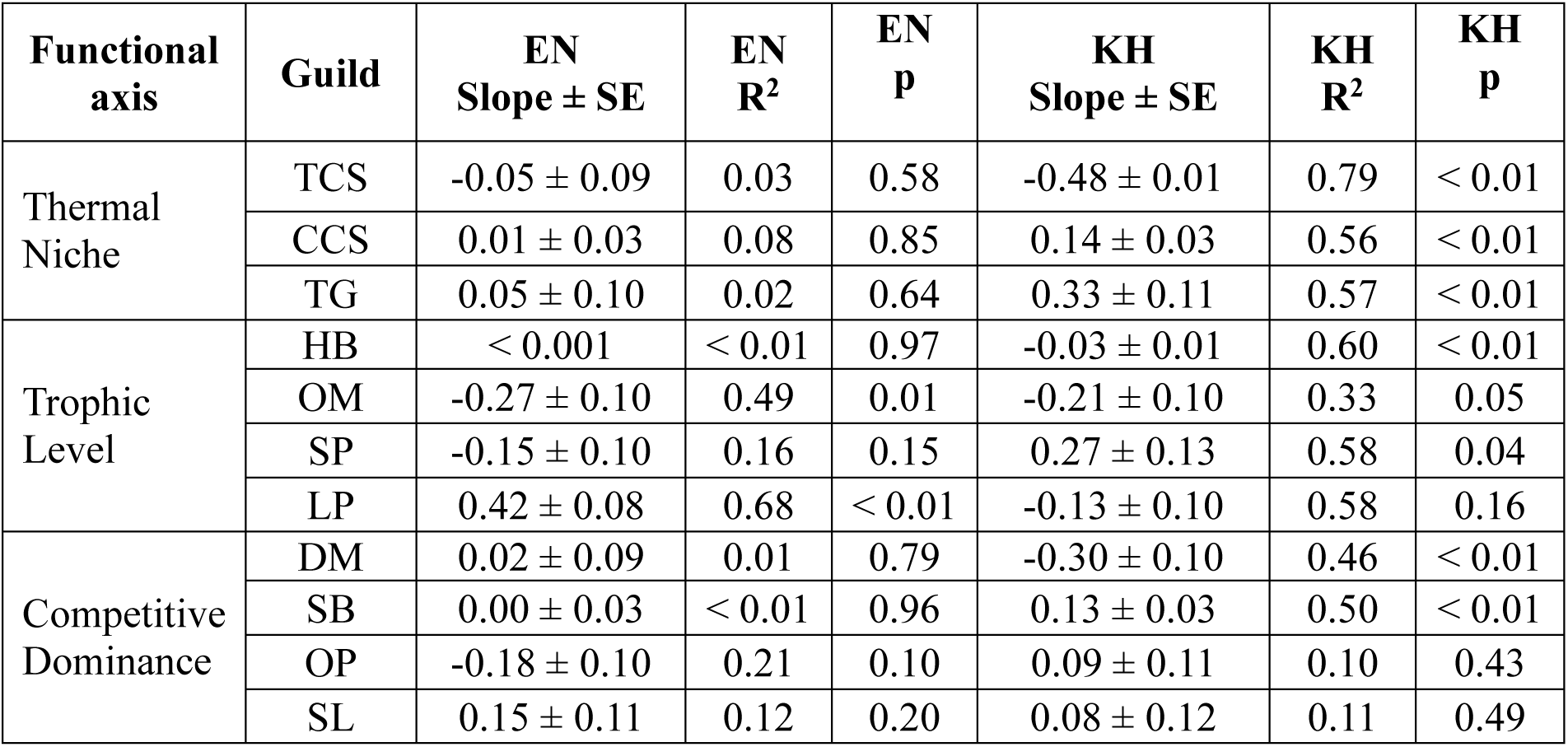
Summary of linear regression between relative abundance and elevation in Eaglenest (EN) and Khupi (KH). Slopes ± SE (per km), p-values and R^2^ statistics are shown for all eleven guilds within the three functional axes (thermal niche, trophic level and competitive dominance. Abbreviations are as follows: TCS = tropical climate specialists, CCS = cold climate specialists, TG = thermal generalists; HB = herbivore, OM = omnivore, SP = small predator, LP = large predator; DM = dominants, SB = subordinates, OP = opportunists, SL = specialists/competition avoiders.

The three thermal guilds exhibited a flat profile in Eaglenest **(Figure 4)**. However, in Khupi, the fractional abundance of cold-climate specialists (slope = 0.14 ± 0.03, R² = 0.56, p < 0.01) and thermal generalists (slope = 0.33 ± 0.11, R² = 0.57, p < 0.01) increased towards higher elevations, at the expense of tropical-climate specialists (slope = −0.48 ± 0.01, R² = 0.79, p < 0.01).

Predatory ants moderately increased in relative abundance while omnivorous ants showed the opposite trend with elevation **(Figure 5)**. However, this trend in predatory ants had contributions from different predation guilds in Eaglenest and Khupi. In Eaglenest, small predators moderately decreased in proportion (slope = −0.15 ± 0.10, R² = 0.16, p = 0.15) while the large ones showed a steeper rise (slope = 0.42 ± 0.08, R² = 0.68, p < 0.01) to yield an overall increase in predatory ants. On the contrary, in Khupi, the large predators showed a marginal decrease with elevation (slope = −0.13 ± 0.10, R² = 0.58, p = 0.16) but the small predators more than compensated for this (slope = 0.27 ± 0.13, R² = 0.58, p = 0.04) resulting in an overall increase in the proportion of predatory ants.

**Figure 5:**
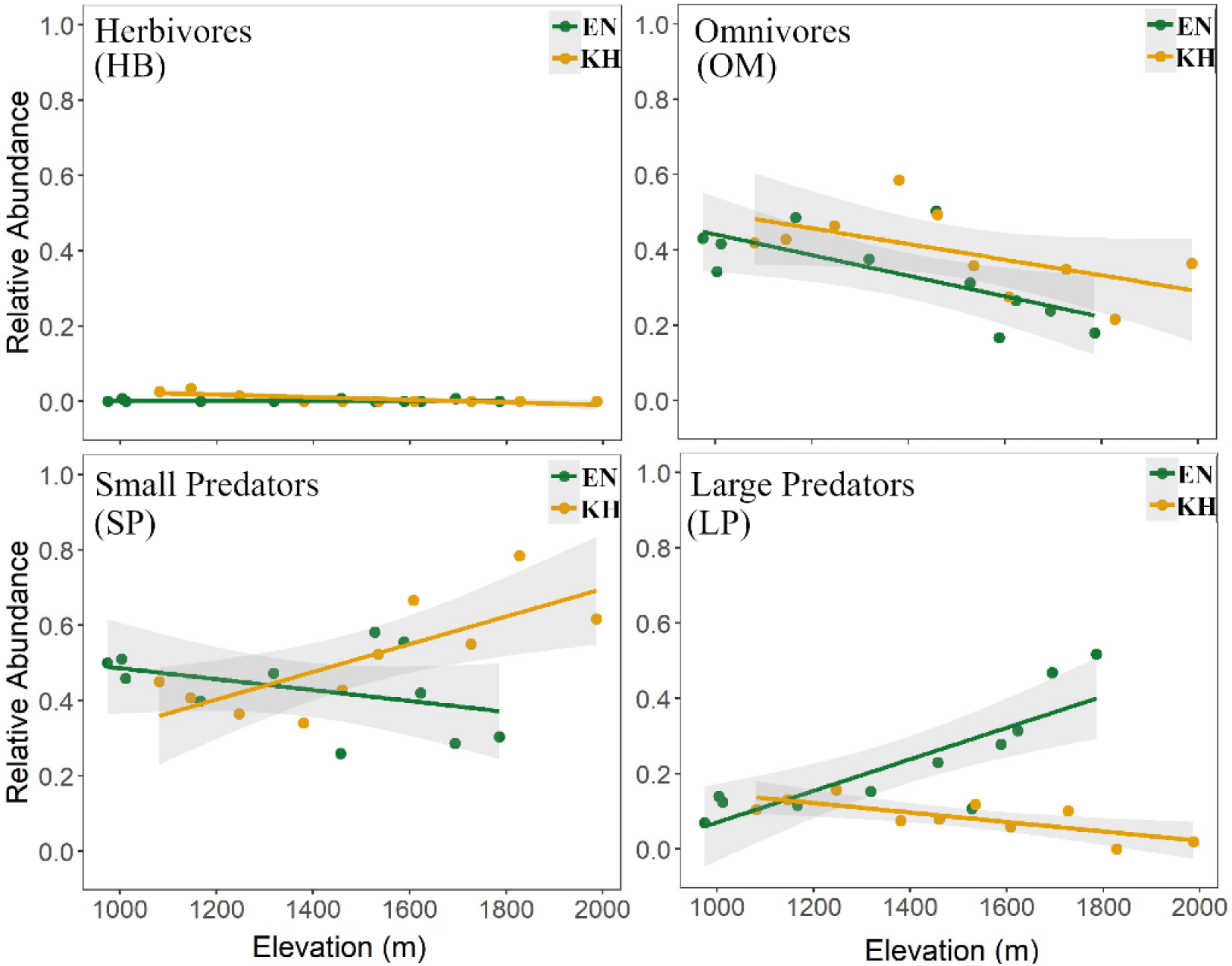
Relative abundance profiles of ant trophic guilds in Eaglenest (EN) and Khupi (KH) along 1000–2000 m span. Each green and yellow solid circle represent an elevational community in Eaglenest and Khupi, respectively. Green and yellow trend lines are the linear fits for the trophic guild profiles with elevation. Abbreviations: TCS = tropical climate specialists, CCS = cold climate specialists, TG = thermal generalists.

The elevational segregation of competitive dominance guilds is weak along Eaglenest **(Figure 6)**. In Khupi, the subordinate guild increases in relative abundance (slope = 0.13 ± 0.03, R² = 0.50, p < 0.01), while the dominant guild shows a declining trend (slope = −0.30 ± 0.10, R² = 0.46, p < 0.01) with elevation.

**Figure 6:**
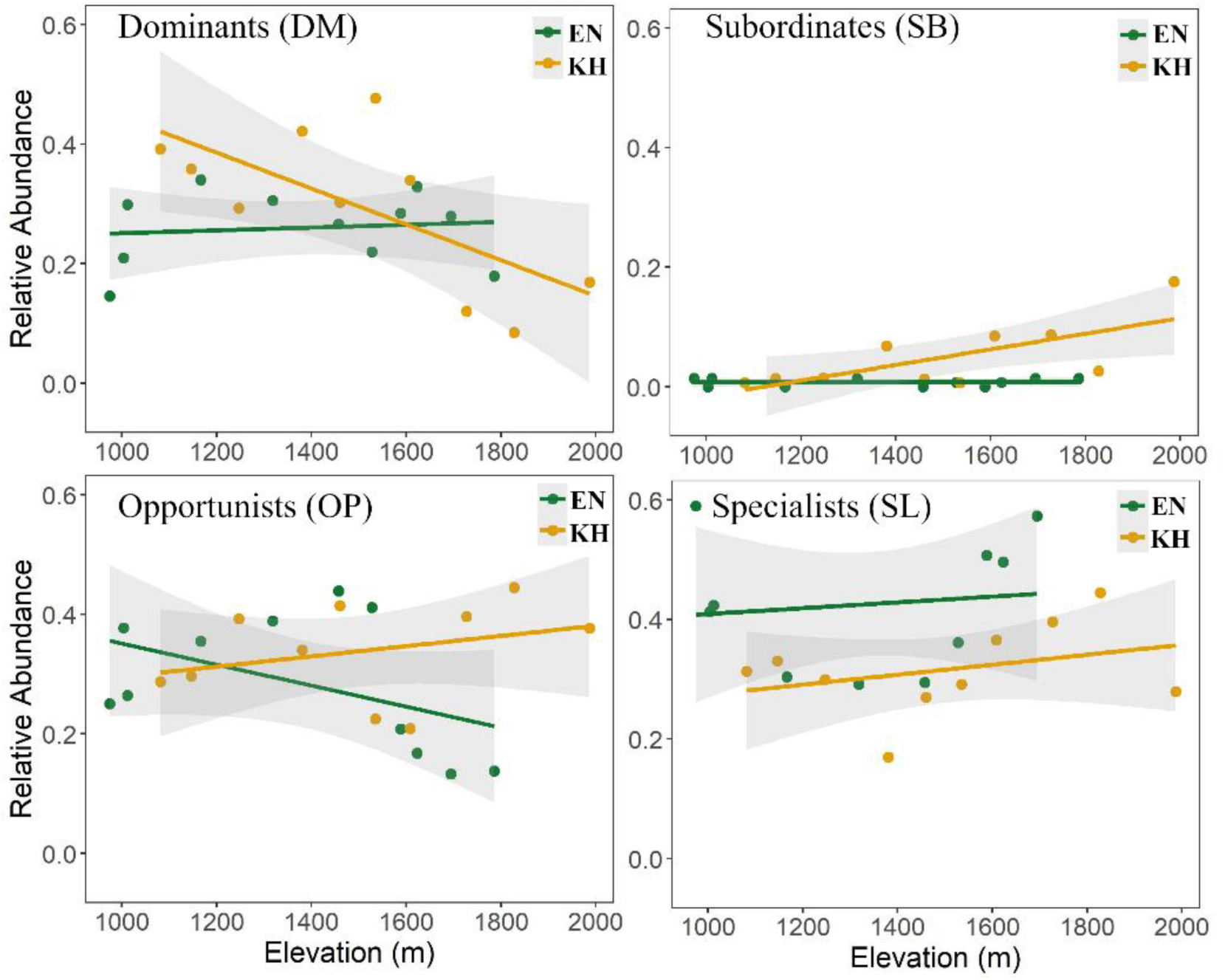
Relative abundance profiles of competitive guilds in ant communities in Eaglenest (EN) and Khupi (KH) along 1000–2000 m span. Each green and yellow solid circle represent an elevational community in Eaglenest and Khupi, respectively. Green and yellow trend lines are the linear fits for the competitive guild profiles with elevation. Abbreviations: TCS = tropical climate specialists, CCS = cold climate specialists, TG = thermal generalists.

The statistical parameters of slope-specific differences in the relative abundance profiles obtained by using an interaction term (for the slopes) are shown in **Table 3**. The composite functional alpha diversity (Rao’s Q) did not change with elevation in both Eaglenest and Khupi **(Figure 7).**

**Figure 7:**
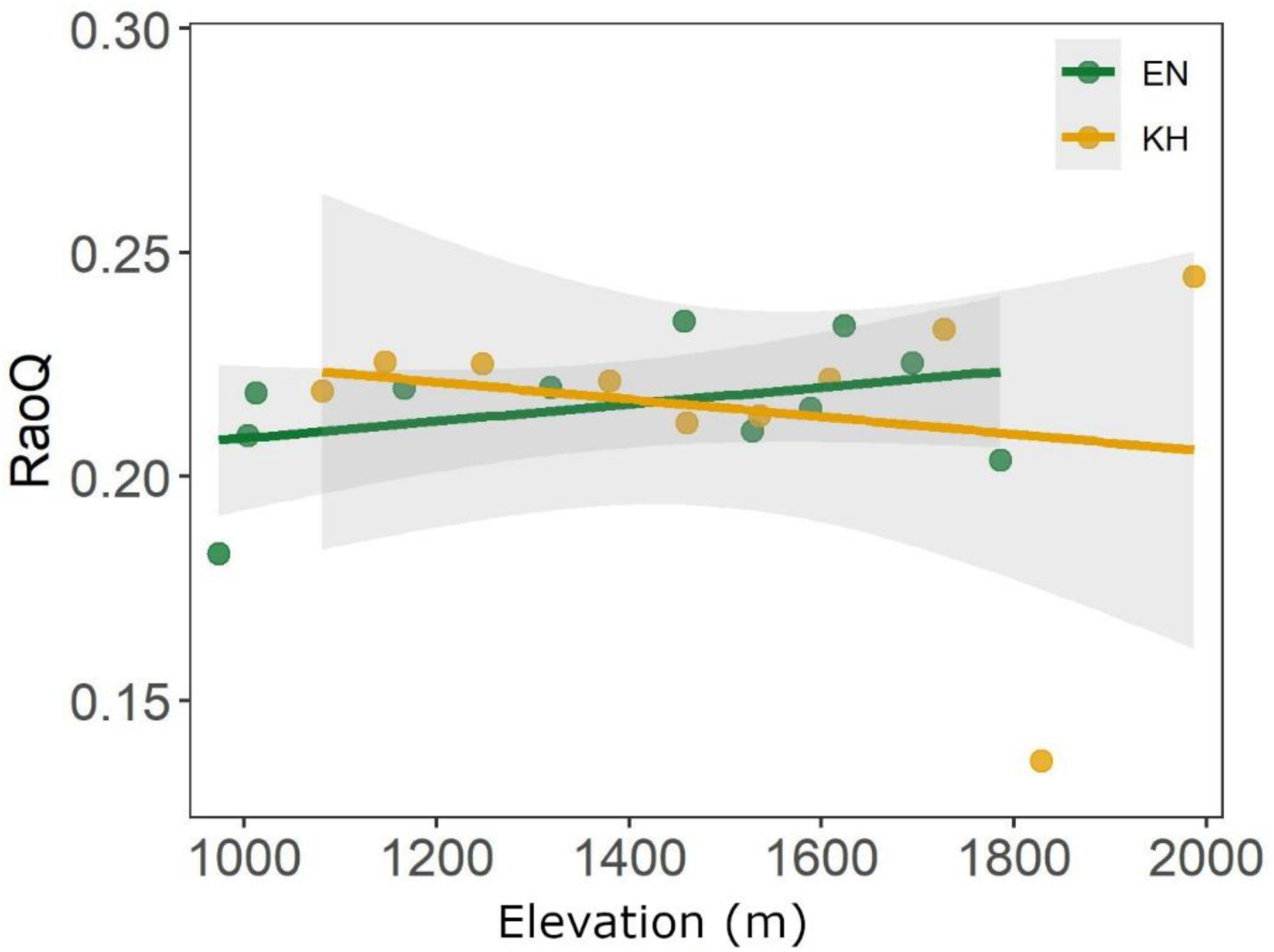
Elevational profiles of functional alpha diversity profiles (Rao’s Q) in Eaglenest (EN) and Khupi (KH) along 1000–2000 m span. Each green and yellow solid circle represent an elevational community in Eaglenest and Khupi, respectively. Green and yellow lines are the linear fits, with 95% confidence intervals.

**Table 3:** Elevation × Transect interaction effects from linear fits examining relative abundance profiles of ant guilds in Eaglenest (EN) and Khupi (KH). The table includes interaction coefficients (β ± SE) and associated p-values for guilds linked to thermal niche, trophic level and competitive dominance. Abbreviations are as follows: TCS = tropical climate specialists, CCS = cold climate specialists, TG = thermal generalists; HB = herbivore, OM = omnivore, SP = small predator, LP = large predator; DM = dominants, SB = subordinates, OP = opportunists, SL = specialists/competition avoiders.

| Functional axis | Guild | Interaction $\beta$ | Interaction p |
| --- | --- | --- | --- |
| Thermal Niche | TCS | $-0.43 \pm 0.14$ | $< 0.01$ |
| | CCS | $0.14 \pm 0.04$ | $< 0.01$ |
| | TG | $0.29 \pm 0.14$ | 0.06 |
| Trophic Level | HB | $-0.03 \pm 0.01$ | $< 0.01$ |
| | OM | $0.07 \pm 0.14$ | 0.64 |
| | SP | $0.43 \pm 0.16$ | $< 0.05$ |
| | LP | $-0.55 \pm 0.12$ | $< 0.01$ |
| Competitive Dominance | DM | $-0.32 \pm 0.13$ | $< 0.05$ |
| | SB | $0.13 \pm 0.04$ | $< 0.01$ |
| | OP | $0.26 \pm 0.15$ | 0.09 |
| | SL | $-0.07 \pm 0.17$ | 0.69 |

Composite functional beta diversity (FβD) increased moderately with elevational distance in both Eaglenest and Khupi **(Figure 8 & Table 4)**. Both turnover and nestedness components of FβD increased with elevational distance, with turnover (mean = 0.32) contributing more than nestedness (mean = 0.045). However, the relationship between composite FβD and its additive components with elevational distance was similar in both Eaglenest and Khupi as inferred from the non-significant interaction coefficients **(Table 4)**.

**Figure 8:**
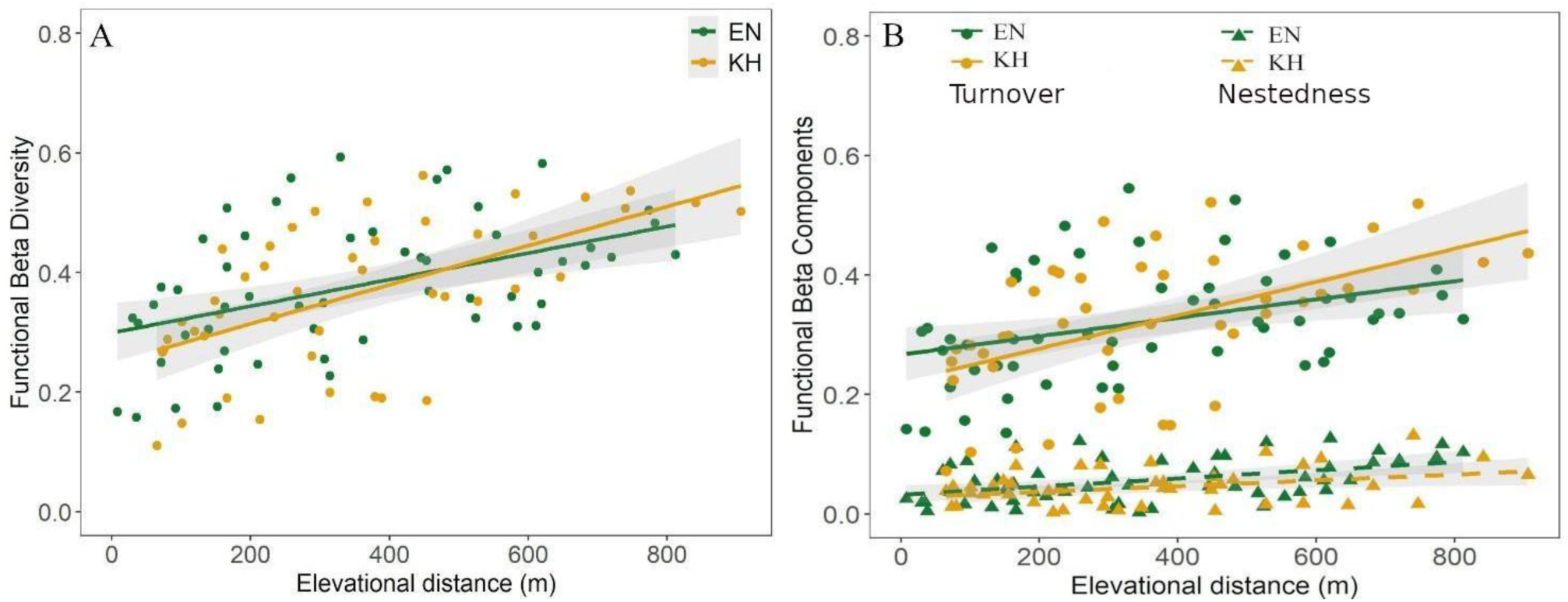
Composite functional beta diversity patterns (A) and its additive components (B) for ant communities in Eaglenest (EN) and Khupi (KH). In panel A, green and yellow solid circles represent total functional beta diversity of elevational communities in Eaglenest and Khupi, respectively. Corresponding solid lines represent the modelled linear relationships between functional beta diversity and elevational distance. Panel B plots the contributions of the two additive components i.e. turnover (circles) and nestedness (triangles), to the observed total functional beta diversity. Green and yellow shapes represent communities values from Eaglenest and Khupi, respectively.

**Table 4:** Summary statistics of total ant functional alpha and beta diversity along with beta components across elevation in Eaglenest (EN) and Khupi (KH). The table reports the estimates of slope ± SE (per km), R^2^ and p-values for functional diversity patterns along each mountain slope. Interaction coefficients (Elevation × Transect) and associated p-values are included to assess whether there are significant differences in elevational patterns between slopes.

| Functional Diversity | EN Slope ± SE | EN $R^2$ | EN p | KH Slope ± SE | KH $R^2$ | KH p | Interaction $\beta$ | Interaction p |
| --- | --- | --- | --- | --- | --- | --- | --- | --- |
| FαD (RaoQ) | 0.019 ± 0.025 | 0.15 | 0.45 | -0.019 ± 0.026 | 0.04 | 0.47 | -0.04 ± 0.04 | 0.30 |
| FβD (Total) | 0.22 ± 0.06 | 0.22 | < 0.01 | 0.33 ± 0.07 | 0.35 | < 0.01 | 0.11 ± 0.09 | 0.24 |
| Functional Turnover | 0.15 ± 0.06 | 0.14 | < 0.01 | 0.28 ± 0.07 | 0.28 | < 0.01 | 0.13 ± 0.09 | 0.15 |
| Functional Nestedness | 0.07 ± 0.02 | 0.20 | < 0.01 | 0.05 ± 0.02 | 0.12 | 0.04 | -0.02 ± 0.46 | 0.46 |

Beta diversity profiles of the individual guilds within each of the three functional axes differed amongst each other and between slopes **(Table 5 & Table 6)**. Individual guilds exhibited steeper profiles in Khupi (7 of the 11 guilds had statistically significant ΔFβD > 10% per km) than in Eaglenest (only 2 guilds).

**Table 5:**
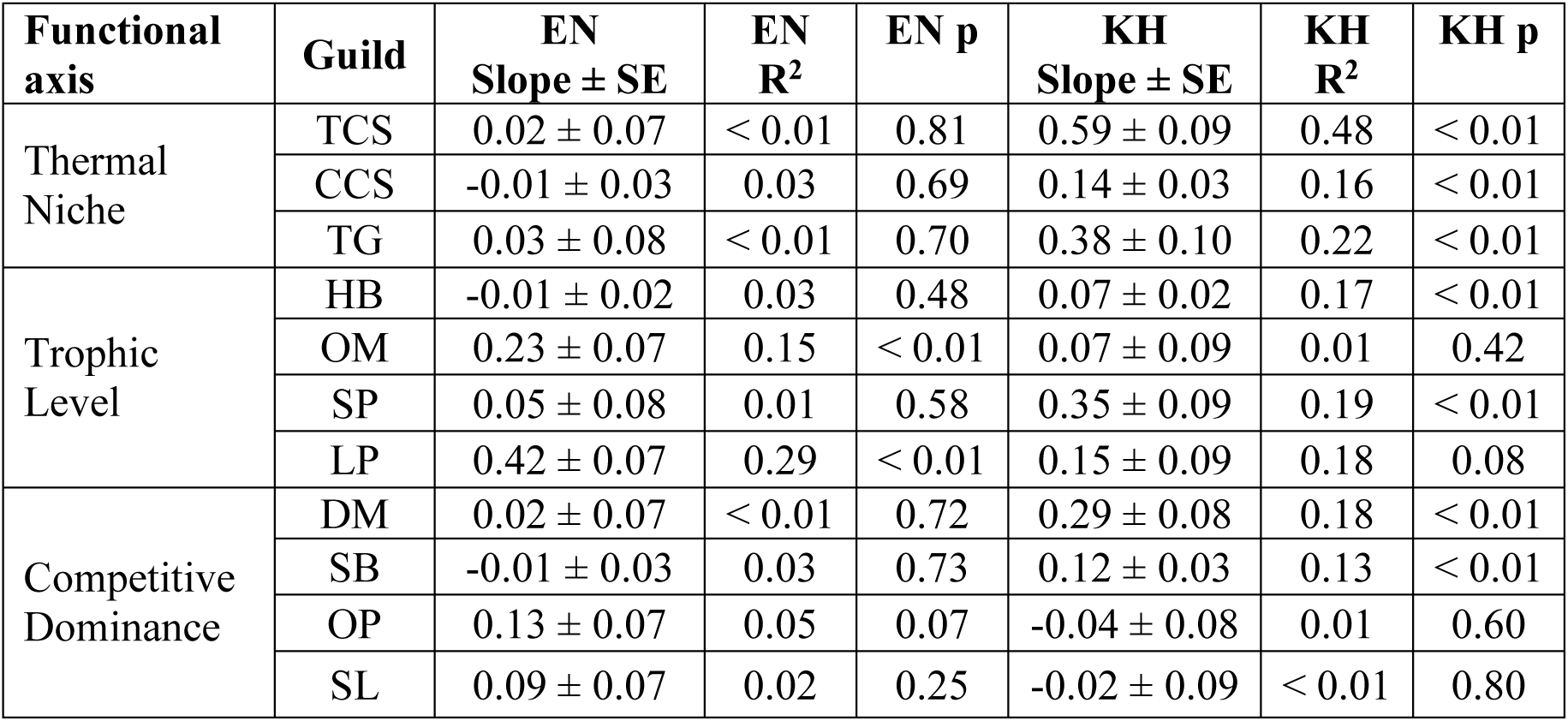
Summary of linear models examining elevational beta diversity profiles of individual functional guilds in Eaglenest (EN) and Khupi (KH). Slopes ± SE (per km), along with R^2^ and p-values are shown for all eleven guilds within each of three functional axes (thermal niche, trophic level and competitive dominance. Abbreviations are as follows: TCS = tropical climate specialists, CCS = cold climate specialists, TG = thermal generalists; HB = herbivore, OM = omnivore, SP = small predator, LP = large predator; DM = dominants, SB = subordinates, OP = opportunists, SL = specialists/competition avoiders.

**Table 6:** Elevation × Transect interaction effects from linear fits examining guild-specific beta diversity profiles in ant communities in Eaglenest (EN) and Khupi (KH). The table includes interaction coefficients (β ± SE) and associated p-values for guilds linked to thermal niche, trophic level and competitive dominance. Abbreviations are as follows: TCS = tropical climate specialists, CCS = cold climate specialists, TG = thermal generalists; HB = herbivore, OM = omnivore, SP = small predator, LP = large predator; DM = dominants, SB = subordinates, OP = opportunists, SL = specialists/competition avoiders.

| Functional axis | Guild | Interaction $\beta$ | Interaction p |
| --- | --- | --- | --- |
| Thermal Niche | TCS | $0.57 \pm 0.11$ | < 0.01 |
| | CCS | $0.15 \pm 0.04$ | < 0.01 |
| | TG | $0.35 \pm 0.12$ | < 0.01 |
| Trophic Level | HB | $0.08 \pm 0.02$ | < 0.01 |
| | OM | $-0.16 \pm 0.11$ | 0.16 |
| | SP | $0.27 \pm 0.12$ | < 0.05 |
| | LP | $-0.27 \pm 0.11$ | < 0.05 |
| Competitive Dominance | DM | $0.26 \pm 0.10$ | 0.01 |
| | SB | $0.13 \pm 0.04$ | < 0.01 |
| | OP | $-0.18 \pm 0.11$ | 0.11 |
| | SL | $-0.11 \pm 0.11$ | 0.35 |

Thermal guilds exhibited flat beta profiles with elevational distance in Eaglenest, while in Khupi they showed increasing trends **(Figure 8)**. Of the three thermal guilds, tropical-climate specialists exhibited the steepest change in Khupi (slope = 0.59 ± 0.09, R² = 0.48, p < 0.01) For trophic guilds, omnivores (slope = 0.23 ± 0.07, R² = 0.15, p < 0.01) and large predators (slope = 0.42 ± 0.07, R² = 0.29, p < 0.01) in Eaglenest show significant increase in FβD with elevational distance. In Khupi, small (slope = 0.35 ± 0.09, R² = 0.19, p < 0.01) and large predators (slope = 0.15 ± 0.09, R² = 0.18, p = 0.08), but not omnivores (slope = 0.07 ± 0.09, R² = 0.01, p = 0.42), exhibited increasing FβD with elevational distance **(Figure 9)**. Eight of eleven guilds exhibited slope-specific differences in Khupi **(Table 5)**.

**Figure 9:**
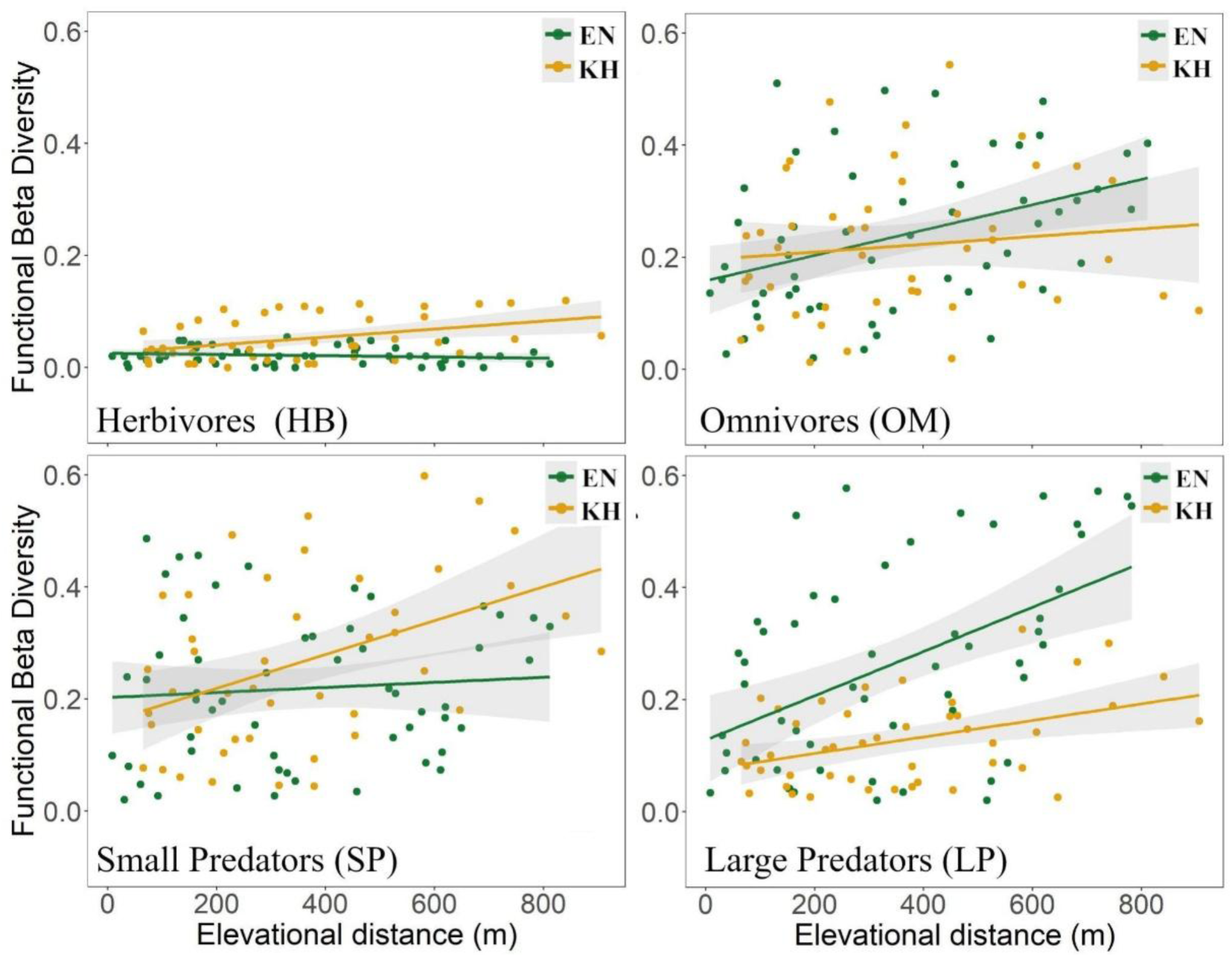
Functional beta diversity profiles of trophic guilds in Eaglenest (EN) and Khupi (KH) along 1000–2000 m span. Each green and yellow solid circle represent an elevational community in Eaglenest and Khupi, respectively. Green and yellow trend lines represent the linear fits showing compositional change within each trophic guild with elevation. Abbreviations: TCS = tropical climate specialists, CCS = cold climate specialists, TG = thermal generalists.

## Discussion

We investigated if differences in local topographic resulted in elevational patterns of functional diversity in Himalayan ant communities. Despite their topographic and associated differences, the two mountain slopes exhibited similar patterns of multivariate functional α-and β-diversity. In contrast, the elevational responses of individual functional guilds differed substantially between the slopes. These findings suggest that similar levels of overall functional diversity can be maintained through alternative guild-level community configuration, with changes in the relative abundances and turnover of individual guilds compensating for one another. Consequently, local topographic heterogeneity may strongly influence the mechanisms of community assembly without necessarily altering the amount of functional space occupied by a community. We discuss these results in detail below.

### Relative Abundance of guilds

Despite being located just 30 km apart, Eaglenest and Khupi exhibited different rank abundance hierarchies of ant thermal guilds **(Figure 3)**. The south-facing slopes of Eaglenest receive higher rainfall and also provide greater opportunity for direct solar warmth even during winter. This may account for the dominance of tropical-climate specialists over thermal generalists in Eaglenest, while the two guilds are co-dominant in north-facing Khupi. Local microclimatic conditions are known to play a significant role in structuring thermal guilds in ant communities (Nascimento et al. 2022), and more generally in arthropods (Khaliq et al., 2025).

There was no difference in the rank hierarchy of trophic guilds between the two slopes. We observed the dominance of small (cryptic) predators on both slopes over other trophic groups. This has been previously reported elsewhere in the tropics, but during the rainy season only (Castro et al., 2020). This may also be related to the method of sampling: Winkler traps are known to favour small-bodied arthropods while pitfalls are biased towards larger ones (Cardoso et al. 2011).

### Elevational profiles of guild relative abundance

In Khupi, tropical-climate specialists reduced substantially towards higher elevations, while thermal generalists and cold-climate specialists dominated at higher elevations; as expected while going from the warmer to cooler climates. However, in Eaglenest, we found weak elevational structuring of thermal guilds along a similar elevational span. This is despite the sharp rank hierarchy among the three guilds. Response of ant communities to temperature can often be complex, with different components of the thermal response (e.g., tolerance and sensitivity) responding to changes at different spatial scales. (Leahy et al., 2022).

Elevational structuring of trophic guilds is often associated with changes in resource availability and quality (Binkenstein et al., 2018), which may be driven by differences in soil composition, for instance, at finer spatial scales. We found differing elevational responses among trophic guilds, with omnivores declining and predatory ants increasing in relative abundance with elevation. Predation is known to dominate at higher elevations in ant communities, especially above the tree line (Guariento & Fiedler, 2021). Even though our sampling was limited to forested elevations well below the tree line, the observed increase in relative abundance of predatory ants with elevation is consistent with patterns reported from three tropical (forested) elevational gradients in Tanzania, Papua New Guinea and Ecuador (Moses et al., 2021). However, Binkenstein et al. (2018) from China and Uhey et al. (2022) from USA report the opposite trend from a subtropical and temperate elevational gradient.

Notably, Uhey et al., (2022) reported habitat-specific differences in the response of predatory ants within the same region, with their dominance increasing at higher elevations in the forested transect but declining in the open habitat transect. These differences were not apparent in a previous study that reported convergence in the diversity and abundance profiles of all arthropods groups along the same transects, which may be in response temperature-moisture balance required for arthropod activity (Uhey et al., 2021). In arid regions, water availability may be the limiting factor at low elevations while temperature may limit diversity at higher elevations, leading to a peak at mid-elevations (Supriya et al., 2019). Hence, broad-scale elevational profiles may mask patterns revealed on segregated communities into ecologically meaningful groups.

We find that the similar elevational profiles for predatory ants between the two mountain slopes obscure underlying divergent responses of predatory communities partitioned by body size. In Eaglenest, the increasing dominance of predators at higher elevations was driven by large predators with no contribution from small predators. On the other hand, small predators dominated the elevational signal in Khupi. A previous study in Eaglenest showed a similar pattern of no-change for small predators but a declining pattern for large predators (Marathe et al., 2021). However, the pattern of declining large predators reported by that study in Eaglenest is consistent with our finding in Khupi. Reduced area at higher elevations in Khupi may explain the dominance of small predators, at the expense of large predators, as we sampled up to 2080 m (ridge at 2100 m). Smaller body size is linked to smaller range sizes in insects and vertebrates, largely due to reduced dispersal abilities (Biedermann, 2003). The dominance of large predators at higher elevations in Eaglenest may be due to our transect terminating at 1850 m i.e. well below the ridge at 3200 m.

In Khupi, we observed a decline in dominant ants and an increase in subordinates at higher elevations, while Eaglenest exhibited no such change. The observed shift between dominant and subordinate groups in Khupi could be explained by the dominance-diversity hypothesis that predicts a positive correlation between dominant ants and diversity of communities (Whittaker, 1965), since we found a monotonic decline in Simpson’s diversity with elevation along the same transect (in prep). Other factors such as reduced carbohydrate availability at higher elevations linked to reduced aggression of dominants along with relatively broader thermal tolerances of subordinates may explain our findings (Boet et al., 2020; Contala et al., 2024; Retana & Cerdá, 2000)

### Multi-guild composite responses of functional diversity with elevation

Even though we found considerable variation in elevational profiles of the relative abundance of functional guilds, we found no change in composite functional α-diversity with elevation in both Khupi and Eaglenest. Studies in subtropical and temperate systems have reported declining patterns of ant functional α-diversity with elevation (Fontanilla et al., 2019; Reymond et al., 2013; Silvestre et al., 2021). Reduction in the prevalence of some functional guilds are offset by increases in other, with elevational communities changing in configuration despite maintaining similar functional space (De Bello et al., 2021). A previous study from the region reported a decline in taxonomic richness within each functional group with elevation along with reduced functional evenness (Maratheet al., 2021). We note that Marathe et al., (2021) does not include a multi-guild framework for each genus and could not directly analyse functional diversity.

As expected, total functional beta diversity increased with elevational distance in both Eaglenest and Khupi. The change in functional composition was largely driven by turnover (88%) rather than nestedness. Studies in ant communities have reported significant contributions from both turnover and nestedness (Bishop et al., 2015; Nunes et al., 2020), with aspect- and season-dependent shifts in their relative contributions (Muluvhahothe et al., 2021; Nunes et al., 2020).

Composite functional beta diversity and its components exhibited similar trends in Eaglenest and Khupi, in contrast to a previous report of aspect-driven differences in composite FβD patterns (Muluvhahothe et al., 2021). Interestingly, despite the similar composite FβD, elevational trends in individual guilds differed substantially between the slopes. We observed that seven out of eleven guilds in Khupi showed a significant trend with elevation while only two guilds (omnivores and large predators) did so in Eaglenest. Several studies have reported that the composite patterns is driven by a few traits or guilds, suggesting the dominance of a subset of evolutionarily labile traits rather than a uniform response across the entire assemblage (Altamirano et al., 2020; Jamoneau et al., 2022; Malumbres-Olarte et al., 2025; Mancini et al., 2019; Ndiribe et al., 2014; Uemori et al., 2022). However, some studies have implicated multiple traits reflecting the trend in the composite pattern (Ding et al., 2019; Read et al., 2014).

Topography, microclimate and seasonality are some factors (other than elevation) that may lead to variability in the observed composite and guild-specific patterns in communities. Microclimatic variation over small elevational distances can parallel vast latitudinal temperature gradients (McNichol et al., 2024), and may filter specific functions along similar macroclimatic extents (Lajoie & Vellend, 2015; Spasojevic et al., 2016). Factors like moisture regimes and topography drive variability in traits like body size and seed weight across slopes (Hishi et al., 2022; Tang et al., 2025). Ant colonies behave like sessile plants (Andersen, 1995), showing more sensitivity to microclimatic filters compared to fauna with greater dispersal abilities (Barton et al., 2024; Heydari et al., 2021). These results highlight the importance of separating macroclimatic and microclimatic factors while investigating functional diversity patterns in ants.

## Acknowledgements

We thank the Forest Department of Arunachal Pradesh for granting research permits for this study; Gendan Marphew, Gore Rana, Bayung and Baiduwa for carrying out the field campaign; the Bugun community for help in the field; Anand Nadathur for partial financial support; IISER Pune for partial financial support and research facilities.

## Conflict of interest

There is no conflict of interest involved in this study

## Ethics approval

Sampling of ants does not require ethics approval as per current regulations

## Author Contributions

Netra Kadambi organised the field campaign, curated the ant samples, did the analysis, surveyed the literature and wrote the first draft of the manuscript. Mansi Mungee partially supervised the analysis and contributed to the manuscript writing. Ramana Athreya contributed to all aspects of the project in a supervisory role.

---

**Figure 8:**
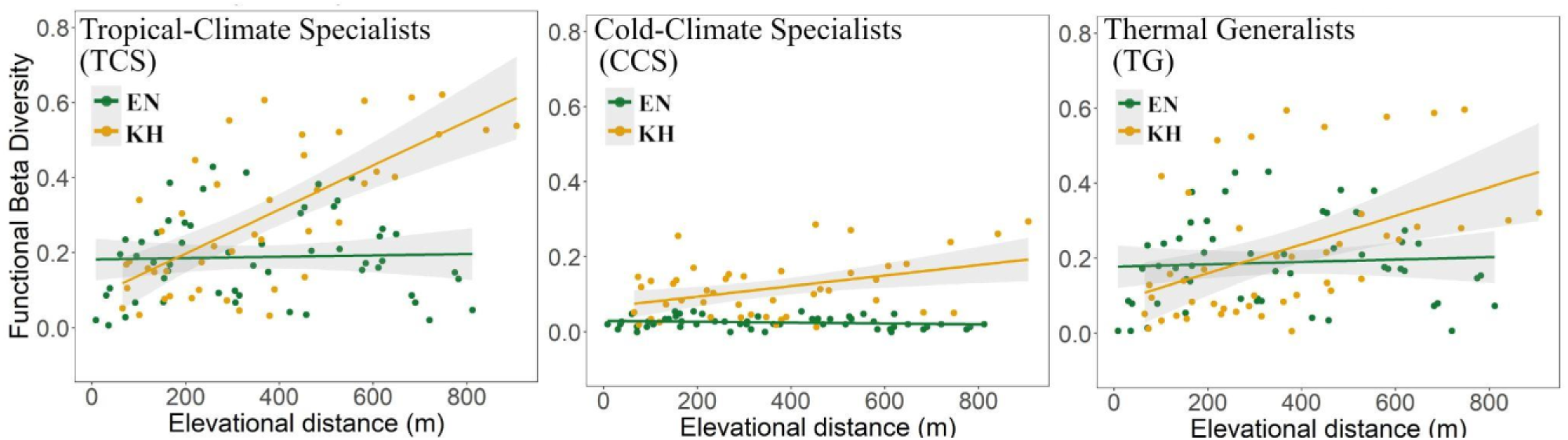
Functional beta diversity profiles of thermal guilds in Eaglenest (EN) and Khupi (KH) along 1000–2000 m span. Each green and yellow solid circle represent an elevational community in Eaglenest and Khupi, respectively. Green and yellow trend lines represent the linear fits showing compositional change within each thermal guild with elevation. Abbreviations: TCS = tropical climate specialists, CCS = cold climate specialists, TG = thermal generalists.

## Notes

### Competing Interest Statement

The authors have declared no competing interest.

## References

Altamirano, T. A., de Zwaan, D. R., Ibarra, J. T., Wilson, S., & Martin, K. (2020). Treeline ecotones shape the distribution of avian species richness and functional diversity in south temperate mountains. Scientific Reports, 10(1), 18428. 10.1038/s41598-020-75470-2

Andersen, A. N. (1995). A Classification of Australian Ant Communities, Based on Functional Groups Which Parallel Plant Life-Forms in Relation to Stress and Disturbance. Journal of Biogeography, 22(1), 15–29. 10.2307/2846070

Anderson, M. J., Crist, T. O., Chase, J. M., Vellend, M., Inouye, B. D., Freestone, A. L., Sanders, N. J., Cornell, H. V., Comita, L. S., Davies, K. F., Harrison, S. P., Kraft, N. J. B., Stegen, J. C., & Swenson, N. G. (2011). Navigating the multiple meanings of β diversity: A roadmap for the practicing ecologist. Ecology Letters, 14(1), 19–28. 10.1111/j.1461-0248.2010.01552.x

Barton, P. S., Evans, M. J., & Lewis, J. (2024). Microhabitats shape ant community structure in a spatially heterogeneous grassy woodland. Ecosphere, 15(4), e4798. 10.1002/ecs2.4798

Basu, P. (1997). Seasonal and Spatial Patterns in Ground Foraging Ants in a Rain Forest in the Western Ghats, India. Biotropica, 29(4), 489–500. 10.1111/j.1744-7429.1997.tb00043.x

Bellwood, D. R., Streit, R. P., Brandl, S. J., & Tebbett, S. B. (2019). The meaning of the term ‘function’ in ecology: A coral reef perspective. Functional Ecology, 33(6), 948–961. 10.1111/1365-2435.13265

Bharti, H., Sharma, Y. P., & Kaur, A. (2009). Seasonal Patterns of Ants (Hymenoptera: Formicidae) in Punjab Shivalik.

Biedermann, R. (2003). Body size and area-incidence relationships: Is there a general pattern? Global Ecology and Biogeography, 12(5), 381–387. 10.1046/j.1466-822X.2003.00048.x

Binkenstein, J., Klein, A.-M., Assmann, T., Buscot, F., Erfmeier, A., Ma, K., Pietsch, K. A., Schmidt, K., Scholten, T., Wubet, T., Bruelheide, H., Schuldt, A., & Staab, M. (2018). Multi-trophic guilds respond differently to changing elevation in a subtropical forest. Ecography, 41(6), 1013–1023. 10.1111/ecog.03086

Bishop, T. R., Robertson, M. P., van Rensburg, B. J., & Parr, C. L. (2015). Contrasting species and functional beta diversity in montane ant assemblages. Journal of Biogeography, 42(9), 1776–1786. 10.1111/jbi.12537

Blaum, N., Mosner, E., Schwager, M., & Jeltsch, F. (2011). How functional is functional? Ecological groupings in terrestrial animal ecology: towards an animal functional type approach. Biodiversity and Conservation, 20(11), 2333–2345. 10.1007/s10531-011-9995-1

Boet, O., Arnan, X., & Retana, J. (2020). The role of environmental vs. biotic filtering in the structure of European ant communities: A matter of trait type and spatial scale. PLOS ONE, 15(2), e0228625. 10.1371/journal.pone.0228625

Castro, F. S. de, Da Silva, P. G., Solar, R., Fernandes, G. W., & Neves, F. de S. (2020). Environmental drivers of taxonomic and functional diversity of ant communities in a tropical mountain. Insect Conservation and Diversity, 13(4), 393–403. 10.1111/icad.12415

Cerda, X., Retana, J., Arnan, X., Angulo, E., & Boulay, R. (2014). Global life trait spectra of resource exploitation in European ants. https://ses.library.usyd.edu.au/handle/2123/10841

Choi, S.-W., & Jang, B.-J. (2022). Effects of elevation and slope on the alpha and beta diversity of ground-dwelling beetles in Mt. Jirisan National Park, South Korea. Journal of Asia-Pacific Entomology, 25(4), 101993. 10.1016/j.aspen.2022.101993

Contala, M.-L., Krapf, P., Steiner, F. M., & Schlick-Steiner, B. C. (2024). Foraging valor linked with aggression: Selection against completely abandoning aggression in the high-elevation ant Tetramorium alpestre? Insect Science, 31(3), 953–970. 10.1111/1744-7917.13263

Davidson, D. W. (1980). Some Consequences of Diffuse Competition in a Desert Ant Community. The American Naturalist, 116(1), 92–105. 10.1086/283613

Dawson, S. K., Carmona, C. P., González-Suárez, M., Jönsson, M., Chichorro, F., Mallen-Cooper, M., Melero, Y., Moor, H., Simaika, J. P., & Duthie, A. B. (2021). The traits of “trait ecologists”: An analysis of the use of trait and functional trait terminology. Ecology and Evolution, 11(23), 16434–16445. 10.1002/ece3.8321

Ding, Y., Zang, R., Lu, X., Huang, J., & Xu, Y. (2019). The effect of environmental filtering on variation in functional diversity along a tropical elevational gradient. Journal of Vegetation Science, 30(5), 973–983. 10.1111/jvs.12786

Drager, K. I., Rivera, M. D., Gibson, J. C., Ruzi, S. A., Hanisch, P. E., Achury, R., & Suarez, A. V. (2023). Testing the predictive value of functional traits in diverse ant communities. Ecology and Evolution, 13(4), e10000. 10.1002/ece3.10000

Feldmeier, S., Schmidt, B. R., Zimmermann, N. E., Veith, M., Ficetola, G. F., & Lötters, S. (2020). Shifting aspect or elevation? The climate change response of ectotherms in a complex mountain topography. Diversity and Distributions, 26(11), 1483–1495. 10.1111/ddi.13146

Fichaux, M., Vleminckx, J., Courtois, E. A., H. C. Delabie, J., Galli, J., Tao, S., Labrière, N., Chave, J., Baraloto, C., & Orivel, J. (2021). Environmental determinants of leaf litter ant community composition along an elevational gradient. Biotropica, 53(1), 97–109. 10.1111/btp.12849

Fontanilla, A. M., Nakamura, A., Xu, Z., Cao, M., Kitching, R. L., Tang, Y., & Burwell, C. J. (2019). Taxonomic and Functional Ant Diversity Along tropical, Subtropical, and Subalpine Elevational Transects in Southwest China. Insects, 10(5), 128. 10.3390/insects10050128

Funk, J. L., Larson, J. E., Ames, G. M., Butterfield, B. J., Cavender-Bares, J., Firn, J., Laughlin, D. C., Sutton-Grier, A. E., Williams, L., & Wright, J. (2017). Revisiting the Holy Grail: Using plant functional traits to understand ecological processes. Biological Reviews, 92(2), 1156–1173. 10.1111/brv.12275

Geres, L. S., Richter, T., Seidl, R., König, S., Chao, A., Chiu, C.-H., Kortmann, M., Mitesser, O., Müller, J., Rothacher, J., Bässler, C., & Seibold, S. (2025). Macro- and microclimate interactively shape species diversity of multiple taxa in mountain landscapes. Ecography, 2025(12), e07984. 10.1002/ecog.07984

Gibb, H., Dunn, R. R., Sanders, N. J., Grossman, B. F., Photakis, M., Abril, S., Agosti, D., Andersen, A. N., Angulo, E., Armbrecht, I., Arnan, X., Baccaro, F. B., Bishop, T. R., Boulay, R., Brühl, C., Castracani, C., Cerda, X., Del Toro, I., Delsinne, T.,…Parr, C. L. (2017). A global database of ant species abundances. Ecology, 98(3), 883–884. 10.1002/ecy.1682

Gilgado, J. D., Rusterholz, H.-P., Braschler, B., Zimmermann, S., Chittaro, Y., & Baur, B. (2022). Six groups of ground-dwelling arthropods show different diversity responses along elevational gradients in the Swiss Alps. PLOS ONE, 17(7), e0271831. 10.1371/journal.pone.0271831

Guariento, E., & Fiedler, K. (2021). Ant Diversity and Community Composition in Alpine Tree Line Ecotones. Insects, 12(3), 219. 10.3390/insects12030219

Heydari, M., Cheraghi, J., Omidipour, R., Mirab-balou, M., & Pothier, D. (2021). Beta diversity of plant community and soil mesofauna along an elevational gradient in a mountainous semi-arid oak forest. Community Ecology, 22(2), 165–176. 10.1007/s42974-021-00046-7

Hishi, T., Kawakami, E., & Katayama, A. (2022). Changes in the abundance and species diversity of Collembola community along with dwarf bamboo density gradient in a mountainous temperate forest of Japan. Applied Soil Ecology, 180, 104606. 10.1016/j.apsoil.2022.104606

Jamoneau, A., Soininen, J., Tison-Rosebery, J., Boutry, S., Budnick, W. R., He, S., Marquié, J., Jyrkänkallio-Mikkola, J., Pajunen, V., Teittinen, A., Tupola, V., Wang, B., Wang, J., Blanco, S., Borrini, A., Cantonati, M., Valente, A. C., Delgado, C., Dörflinger, G.,…Passy, S. I. (2022). Stream diatom biodiversity in islands and continents—A global perspective on effects of area, isolation and environment. Journal of Biogeography, 49(12), 2156–2168. 10.1111/jbi.14482

Jansen, J., Hill, N. A., Dunstan, P. K., Eléaume, M. P., & Johnson, C. R. (2018). Taxonomic Resolution, Functional Traits, and the Influence of Species Groupings on Mapping Antarctic Seafloor Biodiversity. Frontiers in Ecology and Evolution, 6. 10.3389/fevo.2018.00081

König, S., Krauss, J., Classen, A., Hof, C., Prietzel, M., Wagner, C., & Steffan-Dewenter, I. (2024). Micro- and macroclimate interactively shape diversity, niches and traits of Orthoptera communities along elevational gradients. Diversity and Distributions, 30(5), e13810. 10.1111/ddi.13810

Lajoie, G., & Vellend, M. (2015). Understanding context dependence in the contribution of intraspecific variation to community trait–environment matching. Ecology, 96(11), 2912–2922. 10.1890/15-0156.1

Leahy, L., Scheffers, B. R., Williams, S. E., & Andersen, A. N. (2022). Arboreality drives heat tolerance while elevation drives cold tolerance in tropical rainforest ants. Ecology, 103(1), e03549. 10.1002/ecy.3549

Malumbres-Olarte, J., Crespo, L., Cardoso, P., Laizzer, R. L., Mwakisoma, A., Rigal, F., Szűts, T., Pape, T., & Scharff, N. (2025). Within-Habitat β Diversity Increases With Elevation in Tropical Forest Spider Assemblages. African Journal of Ecology, 63(7), e70111. 10.1111/aje.70111

Mancini, M. C. S., Laurindo, R. de S., Hintze, F., Mello, R. de M., & Gregorin, R. (2019). Different bat guilds have distinct functional responses to elevation. Acta Oecologica, 96, 35–42. 10.1016/j.actao.2019.03.004

Marathe, A., Shanker, K., Krishnaswamy, J., & Priyadarsanan, D. R. (2021). Species and functional group composition of ant communities across an elevational gradient in the Eastern Himalaya. Journal of Asia-Pacific Entomology, 24(4), 1244–1250. 10.1016/j.aspen.2021.08.009

McCain, C. M., & Grytnes, J. (2010). Elevational Gradients in Species Richness. In Wiley, Encyclopedia of Life Sciences (1st ed.). Wiley. 10.1002/9780470015902.a0022548

McNichol, B. H., Wang, R., Hefner, A., Helzer, C., McMahon, S. M., & Russo, S. E. (2024). Topography-driven microclimate gradients shape forest structure, diversity, and composition in a temperate refugial forest. Plant-Environment Interactions, 5(3), e10153. 10.1002/pei3.10153

Mlambo, M. C. (2014). Not all traits are ‘functional’: Insights from taxonomy and biodiversity-ecosystem functioning research. Biodiversity and Conservation, 23(3), 781–790. 10.1007/s10531-014-0618-5

Moeslund, J., Arge, L., Bøcher, P., Dalgaard, T., Ejrnæs, R., Odgaard, M., & Svenning, J.-C. (2013). Topographically controlled soil moisture drives plant diversity patterns within grasslands. Biodiversity and Conservation, 22. 10.1007/s10531-013-0442-3

Moses, J., Fayle, T. M., Novotny, V., & Klimes, P. (2021). Elevation and leaf litter interact in determining the structure of ant communities on a tropical mountain. Biotropica, 53(3), 906–919. 10.1111/btp.12914

Muluvhahothe, M. M., Joseph, G. S., Seymour, C. L., Munyai, T. C., & Foord, S. H. (2021). Repeated surveying over 6 years reveals that fine-scale habitat variables are key to tropical mountain ant assemblage composition and functional diversity. Scientific Reports, 11(1), 56. 10.1038/s41598-020-80077-8

Mungee, M., Pandit, R., & Athreya, R. (2021). Taxonomic scale dependency of Bergmann’s patterns: A cross-scale comparison of hawkmoths and birds along a tropical elevational gradient. Journal of Tropical Ecology, 37(6), 302–312. 10.1017/S0266467421000432

Munyai, T. C., & Foord, S. H. (2012). Ants on a mountain: Spatial, environmental and habitat associations along an altitudinal transect in a centre of endemism. Journal of Insect Conservation, 16(5), 677–695. 10.1007/s10841-011-9449-9

Myers, N., Mittermeier, R. A., Mittermeier, C. G., Da Fonseca, G. A. B., & Kent, J. (2000). Biodiversity hotspots for conservation priorities. Nature, 403(6772), 853–858. 10.1038/35002501

Nascimento, G., Câmara, T., & Arnan, X. (2022). Critical thermal limits in ants and their implications under climate change. Biological Reviews, 97(4), 1287–1305. 10.1111/brv.12843

Ndiribe, C., Pellissier, L., Dubuis, A., Vittoz, P., Salamin, N., & Guisan, A. (2014). Plant functional and phylogenetic turnover correlate with climate and land use in the Western Swiss Alps. Journal of Plant Ecology, 7(5), 439–450. 10.1093/jpe/rtt064

Nunes, C. A., Castro, F. S., Brant, H. S. C., Powell, S., Solar, R., Fernandes, G. W., & Neves, F. S. (2020). High Temporal Beta Diversity in an Ant Metacommunity, With Increasing Temporal Functional Replacement Along the Elevational Gradient. Frontiers in Ecology and Evolution, 8. 10.3389/fevo.2020.571439

Palacio, F. X., Ottaviani, G., Mammola, S., Graco-Roza, C., de Bello, F., & Carmona, C. P. (2025). Integrating intraspecific trait variability in functional diversity: An overview of methods and a guide for ecologists. Ecological Monographs, 95(2), e70024. 10.1002/ecm.70024

Pérez-Sánchez, A. J., Schibalski, A., Schröder, B., Klimek, S., & Dauber, J. (2023). Local and landscape environmental heterogeneity drive ant community structure in temperate seminatural upland grasslands. Ecology and Evolution, 13(3), e9889. 10.1002/ece3.9889

Perrigo, A., Hoorn, C., & Antonelli, A. (2019). Why mountains matter for biodiversity. Journal of Biogeography | Wiley Online Library. https://onlinelibrary.wiley.com/doi/full/10.1111/jbi.13731

Potter, K. A., Arthur Woods, H., & Pincebourde, S. (2013). Microclimatic challenges in global change biology. Global Change Biology, 19(10), 2932–2939. 10.1111/gcb.12257

Raffard, A., Lecerf, A., Cote, J., Buoro, M., Lassus, R., & Cucherousset, J. (2017). The functional syndrome: Linking individual trait variability to ecosystem functioning. Proceedings of the Royal Society B: Biological Sciences, 284(1868), 20171893. 10.1098/rspb.2017.1893

Rahbek, C., Borregard, M., Antonelli, A., Colwell, R. K., Holt, B. G., Nogues-Bravo, D., Rasmussen, C. M. Ø., Richardson, K., Rosing, M. T., Whittaker, R. J., & Fjeldså, J. (2019). Building mountain biodiversity: Geological and evolutionary processes. Science. https://www.science.org/doi/10.1126/science.aax0151

Read, Q. D., Moorhead, L. C., Swenson, N. G., Bailey, J. K., & Sanders, N. J. (2014). Convergent effects of elevation on functional leaf traits within and among species. Functional Ecology, 28(1), 37–45. 10.1111/1365-2435.12162

Retana, J., & Cerdá, X. (2000). Patterns of diversity and composition of Mediterranean ground ant communities tracking spatial and temporal variability in the thermal environment. Oecologia, 123(3), 436–444. 10.1007/s004420051031

Reymond, A., Purcell, J., Cherix, D., Guisan, A., & Pellissier, L. (2013). Functional diversity decreases with temperature in high elevation ant fauna. Ecological Entomology, 38(4), 364–373. 10.1111/een.12027

Rubio, V. E., & Swenson, N. G. (2024). On functional groups and forest dynamics. Trends in Ecology & Evolution, 39(1), 23–30. 10.1016/j.tree.2023.08.008

Schowalter, T. D., Presley, S. J., & Willig, M. R. (2025). Variation in biodiversity and abundance of functional groups of arthropods along a tropical elevational gradient in Puerto Rico. Biotropica, 57(1), e13412. 10.1111/btp.13412

Silva, P. S. D., Bieber, A. G. D., Corrêa, M. M., & Leal, I. R. (2011). Do leaf-litter attributes affect the richness of leaf-litter ants? Neotropical Entomology, 40, 542–547. 10.1590/S1519-566X2011000500004

Silvestre, M., Carmona, C. P., Azcárate, F. M., & Seoane, J. (2021). Diverging facets of grassland ant diversity along a Mediterranean elevational gradient. Ecological Entomology, 46(6), 1301–1314. 10.1111/een.13077

Spasojevic, M. J., & Suding, K. N. (2012). Inferring community assembly mechanisms from functional diversity patterns: The importance of multiple assembly processes. Journal of Ecology, 100(3), 652–661. 10.1111/j.1365-2745.2011.01945.x

Spasojevic, M. J., Turner, B. L., & Myers, J. A. (2016). When does intraspecific trait variation contribute to functional beta-diversity? Journal of Ecology, 104(2), 487–496. 10.1111/1365-2745.12518

Stein, A., Gerstner, K., & Kreft, H. (2014). Environmental heterogeneity as a universal driver of species richness across taxa, biomes and spatial scales. Ecology Letters, 17(7), 866–880. 10.1111/ele.12277

Sundqvist, M. K., Sanders, N. J., & Wardle, D. A. (2013). Community and Ecosystem Responses to Elevational Gradients: Processes, Mechanisms, and Insights for Global Change. Annual Review of Ecology, Evolution, and Systematics, 44(Volume 44, 2013), 261–280. 10.1146/annurev-ecolsys-110512-135750

Supriya, K., Moreau, C. S., Sam, K., & Price, T. D. (2019). Analysis of tropical and temperate elevational gradients in arthropod abundance. Frontiers of Biogeography, 11(2). 10.21425/F5FBG43104

Swenson, N. G. (2011). The role of evolutionary processes in producing biodiversity patterns, and the interrelationships between taxonomic, functional and phylogenetic biodiversity. American Journal of Botany, 98(3), 472–480. 10.3732/ajb.1000289

Tang, X., Liu, Z., Liu, F., Cheng, Y., Yu, T., Li, X., Qin, Q., & Zhang, F. (2025). Landscape Patterns Drive Functional Diversity of Macroinvertebrate Communities Along the Elevation Gradient in the Chishui River. Biology, 14(9), 1149. 10.3390/biology14091149

Theron, K. J., Pryke, J. S., & Samways, M. J. (2022). Maintaining functional connectivity in grassland corridors between plantation forests promotes high-quality habitat and conserves range restricted grasshoppers. Landscape Ecology, 37(8), 2081–2097. 10.1007/s10980-022-01471-3

Uemori, K., Mita, T., & Hishi, T. (2022). Differences in functional trait responses to elevation among feeding guilds of Aculeata community. Ecology and Evolution, 12(8), e9171. 10.1002/ece3.9171

Uhey, D. A., Bowker, M. A., Haubensak, K. A., Auty, D., Vissa, S., & Hofstetter, R. W. (2022). Habitat Type Affects Elevational Patterns in Ground-dwelling Arthropod Communities. Journal of Insect Science, 22(4), 9. 10.1093/jisesa/ieac046

Violle, C., Enquist, B. J., McGill, B. J., Jiang, L., Albert, C. H., Hulshof, C., Jung, V., & Messier, J. (2012). The return of the variance: Intraspecific variability in community ecology. Trends in Ecology & Evolution, 27(4), 244–252. 10.1016/j.tree.2011.11.014

Violle, C., Navas, M.-L., Vile, D., Kazakou, E., Fortunel, C., Hummel, I., & Garnier, E. (2007). Let the concept of trait be functional! Oikos, 116(5), 882–892. 10.1111/j.0030-1299.2007.15559.x

